# Experimental multi-species microbial (co)evolution results in local maladaptation

**DOI:** 10.1101/2020.04.26.062067

**Authors:** Meaghan Castledine, Daniel Padfield, Angus Buckling

## Abstract

Interspecific coevolutionary interactions can result in rapid biotic adaptation, but most studies have focused only on species pairs. Here, we (co)evolved five microbial species in replicate polycultures and monocultures and quantified local adaptation. Specifically, growth rate assays were used to determine adaptations of each species’ populations to (1) the presence of the other four species in general and (2) sympatric vs allopatric communities. We found no evidence for general biotic adaptation: ancestral, polyculture- and monoculture-evolved populations did not have significantly different growth rates when measured within communities. However, 4/5 species’ growth rates were significantly lower within the community they evolved in. This “local maladaptation” suggests that species evolved increased competitive interactions to sympatric species’ populations. This increased competition did not affect community stability or productivity. Our results suggest that (co)evolution within communities can increase competitive interactions that are specific to (co)evolved community members.

## Introduction

(Co)evolutionary interactions between different species can result in rapid adaptation to biotic conditions (Savolainen *et al*. 2013; Briscoe Runquist *et al*. 2020). The majority of the work on local adaptation has focused on pairwise interactions (Brockhurst & Koskella 2013) with few studies focusing on interactions between multiple community members. While a pairwise perspective may be appropriate for many specialised host-parasite interactions (Poulin & Morand 2000), it will be less so for competitive and mutualistic interactions which often involve complex interaction networks (Rohr *et al*. 2014; Levine *et al*. 2017). Here, we investigated biotic adaptation within a multi-species microbial community dominated by competitive interactions.

Biotic adaptation can vary in specificity. First, adaptation may occur simply to the presence (versus absence) of an interacting species (Kawecki & Ebert 2004; Savolainen *et al*. 2013). Such biotic adaptation can constrain adaptation to other aspects of the abiotic or biotic environment as a result of trade-offs or changes in effective population sizes (Hereford 2009; Lawrence *et al*. 2012; Gómez & Buckling 2013; Friman & Buckling 2014; Hall *et al*. 2018; Briscoe Runquist *et al*. 2020). Examples of abiotic – biotic trade-offs have previously been observed in multi-species communities and the abiotic environment (Lawrence *et al*. 2012; Briscoe Runquist *et al*. 2020). Second, divergence in adaptations can also occur between different populations of the same species which is typically referred to as “local adaptation” (Kawecki & Ebert 2004; Savolainen *et al*. 2013). Much of this work has focused on host-parasite interactions where either parasites, hosts (or neither) can show a tendency to be locally adapted, depending on the study system (Morand *et al*. 1996; Thompson 2002; Morgan *et al*. 2005; Greischar & Koskella 2007; Brady *et al*. 2019).

Coevolution within competitive multi-species communities may result in many of the interacting species being locally adapted. Competitive interactions can lead to divergent selection and character displacement to reduce the extent of interspecific competition (Pacala & Roughgarden 1982; Schluter 2000; Dayan & Simberloff 2005; Leimar *et al*. 2008; Pfennig & Pfennig 2010; Vellend 2010; Burgess *et al*. 2013; terHorst *et al*. 2018). Assuming divergent outcomes between replicate coevolving communities (Stenseth & Smith 1984; terHorst *et al*. 2018), populations are expected to be locally adapted to their community because they are likely to experience less niche overlap with species they have coevolved with. This greater niche partitioning may in turn result in greater community stability (i.e. species are less prone to extinction) and more productive communities as a consequence of more efficient resource use (Dayan & Simberloff 2005; Pfennig & Pfennig 2010; Coyte *et al*. 2015; Edwards *et al*. 2018; terHorst *et al*. 2018). However, if species compete for essential resources, there may be trait convergence and niche overlap (Abrams 1987; Fox & Vasseur 2008; Hart *et al*. 2019) and species may also evolve to increase the direct harm they cause competitors. These outcomes could potentially lead to some species being locally adapted or locally maladapted, as is the case with antagonistic interactions such as between host and parasite pairs (Kawecki & Ebert 2004; Greischar & Koskella 2007; Savolainen *et al*. 2013).

The primary aim of this study was to determine whether or not there is a tendency for populations to become locally adapted following evolution in multi-species communities: i.e. if they are better adapted to the community they evolved in compared to novel communities. To this end, we experimentally (co)evolved soil bacteria species, propagated as replicate polycultures in nutrient media in which they stably coexist (Padfield *et al*. bioRxiv). We compared the magnitude of local adaptation to adaptation to biotic versus abiotic conditions *per se* by also evolving the species in monoculture (Lawrence *et al*. 2012; Briscoe Runquist *et al*. 2020). Finally, we determined if any (co)evolved interactions affected community level properties: stability (the ability of species to invade from rare (Chesson 2018)) and community productivity.

## Materials and methods

### Experimental evolution and clonal isolation

Species isolates were originally obtained from soil and identified as: *Achromobacter* sp. (A), *Ochrobactrum* sp. (O), *Pseudomonas* sp. (P), *Stenotrophomonas* sp. (S) and *Variovorax* sp. (V) (Padfield *et al*., bioRxiv). Each species were grown from one colony (clone) in isolation for two days in 6mL growth media (1/64 Tryptone Soy Broth (TSB); diluted with demineralised H2O), static, at 28°C in 25mL glass microcosms with loosened plastic lids. Twelve replicate polyculture (all species) and monoculture lines (twelve replicates per species) were set-up using 20μL inoculum of each species’ culture into fresh growth media. Inoculated abundance (colony forming units: CFUs) of each species were estimated approximately from optical densities (OD_600_; wavelength 600nm) after two days growth (equations for converting OD_600_ to CFU/μL described by Padfield et al. (bioRxiv)) as: A = 3.5 *x* 10^7^; O = 1.0 *x* 10^8^; P = 1.3 *x* 10^7^; S = 3.6 *x* 10^8^; V = 1.0 *x* 10^8^. Serial 100-fold dilutions (60μL culture into 6mL growth media) took place every week for a total of ten weeks. Culture samples were cryogenically frozen at −80°C in glycerol (final concentration: 25%) every second transfer. After ten weeks, cultures were plated onto KB (King’s Medium B) agar and incubated for two days at 28°C. Six clones of each species per replicate were isolated and grown overnight in growth media before being frozen at −80°C in glycerol. Prior to being used in experimental trials, clones were grown for two days in growth media at 28°C on an orbital shaker at 180 r.p.m. to achieve high cell densities. For all assays, only one clone of each species was used to control for any effects of genetic variation on growth rate (Hughes *et al*. 2008). Where multiple morphotypes had evolved, only the most common morph was isolated.

### Growth rate assays

Using isolated clones, we conducted a series of growth rate assays to assess adaptation (Figure 1). Each polyculture line was grouped into blocks with one monoculture line of each species (i.e. one polyculture line was grouped with one monoculture each of A, O, P, S and V). To measure whether populations adapted to the presence of the other species, communities were re-assembled from isolated clones of the same polyculture line. These growth rates were contrasted to ancestral and monoculture-evolved populations of each species when in a community in which all other species had evolved in polyculture. To assess abiotic adaptation, we measured the growth rates of these same clones in isolation. To assess local adaptation between coevolved communities, the twelve polyculture lines were grouped into six pairs, and the growth rate of each clone was measured in a sympatric and a paired allopatric community.

**Figure 1.**
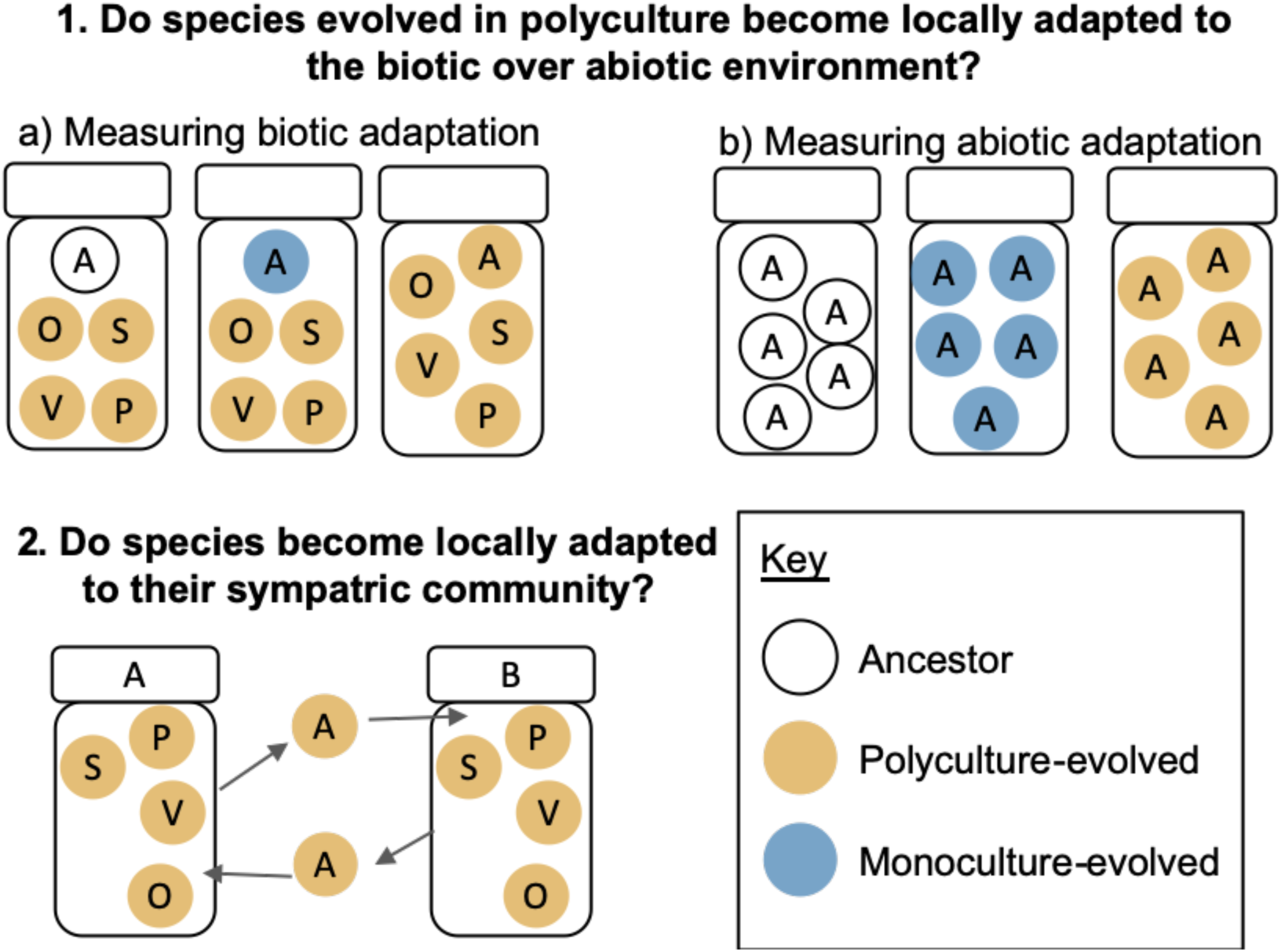
Experimental set-up to test adaptation of each species populations. 1. Ancestral, polyculture-evolved and monoculture-evolved growth rates were compared in communities and in isolation to measure (a) biotic and (b) abiotic adaptation respectively. 2. Growth rates were compared to when species’ populations were in sympatric versus paired allopatric communities. Manipulation of only species A is shown, but all species were exposed to all treatments.

Overall, this established a total of 7 treatments with 12 replicates per species and treatment (Figure 1). The culture conditions of all treatments were established as described for the initial experimental evolution set up above, but with approximately 10^6^ CFUs per species inoculated into fresh microcosms. After one week, culture samples were frozen and then plated onto KB agar. Population densities (CFU/mL) were calculated by counting the number of CFUs after two days of growth at 28°C. Growth rate was calculated using estimated Malthusian parameters, *m* = *ln(N*_*1*_ */ N*_*0*_) / *t* where *N*_*1*_ is the final density, *N*_*0*_ is starting density and *t* is the time given for growth (Lenski *et al*. 1991). As *t* is equal across experiments (1 week), this equation simplifies to *m* = *ln(N*_*1*_ */ N*_*0*_).

### Community stability

To assess whether evolution impacted stability, we invaded monoculture- and polyculture-evolved populations from rare (100-fold lower density to the surrounding community) into monoculture- and polyculture-based communities respectively. All conditions were consistent with the previous design of the 1:1 experiment with approximately ∼10^6^ CFU per resident species and approximately ∼10^4^ CFUs of the invading species. Relative invader fitness of the invading species (*m*_inv_) was calculated as the ratio of estimated Malthusian parameters, *m*_inv_:*m*_community_ where *m*_community_ is the density change of the other four clonal populations combined.

### Community productivity

Additionally, we assessed the effect of evolution on community productivity. Here, we used the data collected from the growth rate assays where polyculture-based communities had been assembled from sympatrically evolved clones, or monoculture-based communities (from clones within the same block) at approximately equal starting densities. We additionally assembled twelve communities randomly from clones of allopatric polyculture lines with clonal populations at approximately equal starting densities (∼10^6^ CFUs per species). One clone per species and no two clones from the same polyculture line were used for each replicate and no clone was used in more than one replicate. Community productivity was defined as the total CFU density change (*ln(N*_*1*_ */ N*_*0*_)) within each community.

### Statistical analyses

Biotic adaptation was contrasted between ancestral, polyculture- and monoculture-evolved populations growing in communities using linear mixed effects models. Abiotic adaptation between ancestral, monoculture- and polyculture-evolved clones of each species was assessed in a linear model. In separate models (for each experimental question), growth rate (*m*) was tested against interacting fixed effects of evolutionary history (ancestor and polyculture- or monoculture-evolved population) and species identity. For the mixed effects model testing biotic adaptation, a random effect of “block” was included to account for which of the assembled communities growth rates had been measured. Local adaptation was tested using a linear mixed effects model in which growth rate was tested against community type (sympatric/allopatric) with species identity as an interacting factor. A random effect of “clone” was included to account for the paired experimental design.

We estimated the ability of each species’ population invade from rare by conducting one sample t-tests on each focal species within each combination with the null hypothesis being that relative invader fitness is equal to 1. A relative invader fitness greater than 1 would indicate that the invading species has a higher growth rate than the other community members, thus its fitness being negatively frequency dependent. This resulted in 10 statistical tests and p-values were corrected by the false discovery rate (*fdr*) method (Benjamini & Hochberg 1995). Additionally, the relative invader fitness of each species’ population was contrasted between community assemblies using a linear model where relative invader fitness was tested against interacting fixed effects of community assembly (monoculture- or polyculture-based) and species identity. Next, the effect of evolution on community productivity was assessed using a linear model in which productivity was analysed against community composition (polyculture [sympatric or allopatric] and monoculture). To compare the structure of these communities (polyculture-sympatric, polyculture-allopatric or monoculture) and the contribution of each species to productivity, we contrasted species proportions between communities. In a general linear model with a quasibinomial error structure (to account for overdispersion), species’ proportions were tested against interacting fixed effects of community assembly (monoculture- or polyculture-based) and species identity.

All data was analysed in R (v 3.5.1) (Team 2013) and all plots were made using the R package ‘*ggplot2*’ (Wickham 2016). Four treatment replicates which became contaminated were removed from analyses (Table S1). Linear mixed effect models were fitted using the R package ‘*lme4*’ (Bates *et al*. 2014). Model simplification was conducted using likelihood ratio tests. Tukey’s post-hoc multiple comparison tests were done on the most parsimonious model using the R package ‘*emmeans*’ (Lenth 2018). In the majority of mixed-effect models fitted, we encountered a singular-fitting issue in which parameter estimates for the random effects could not be estimated. Here, we retained the random effect in the model as they allow for appropriate calculation of degrees of freedom.

## Results

### Biotic and abiotic adaptation

We first considered whether populations evolved in polyculture had adapted to the presence of the other community members in general. To this end, we compared growth rates of clones of ancestral, monoculture- and polyculture-evolved populations within polyculture-evolved communities of the other four species (Figure 1). We did not find evidence of significant biotic adaptation as shown by a non-interacting (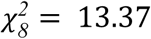, p = 0.072), and a non-significant independent effect of evolutionary history (i.e. whether clones were ancestral, monoculture- or polyculture-evolved; 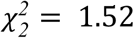, p = 0.467; Figure 2a). Growth rates significantly differed between species (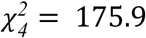, p < 0.001; Figure 2a).

**Figure 2.**
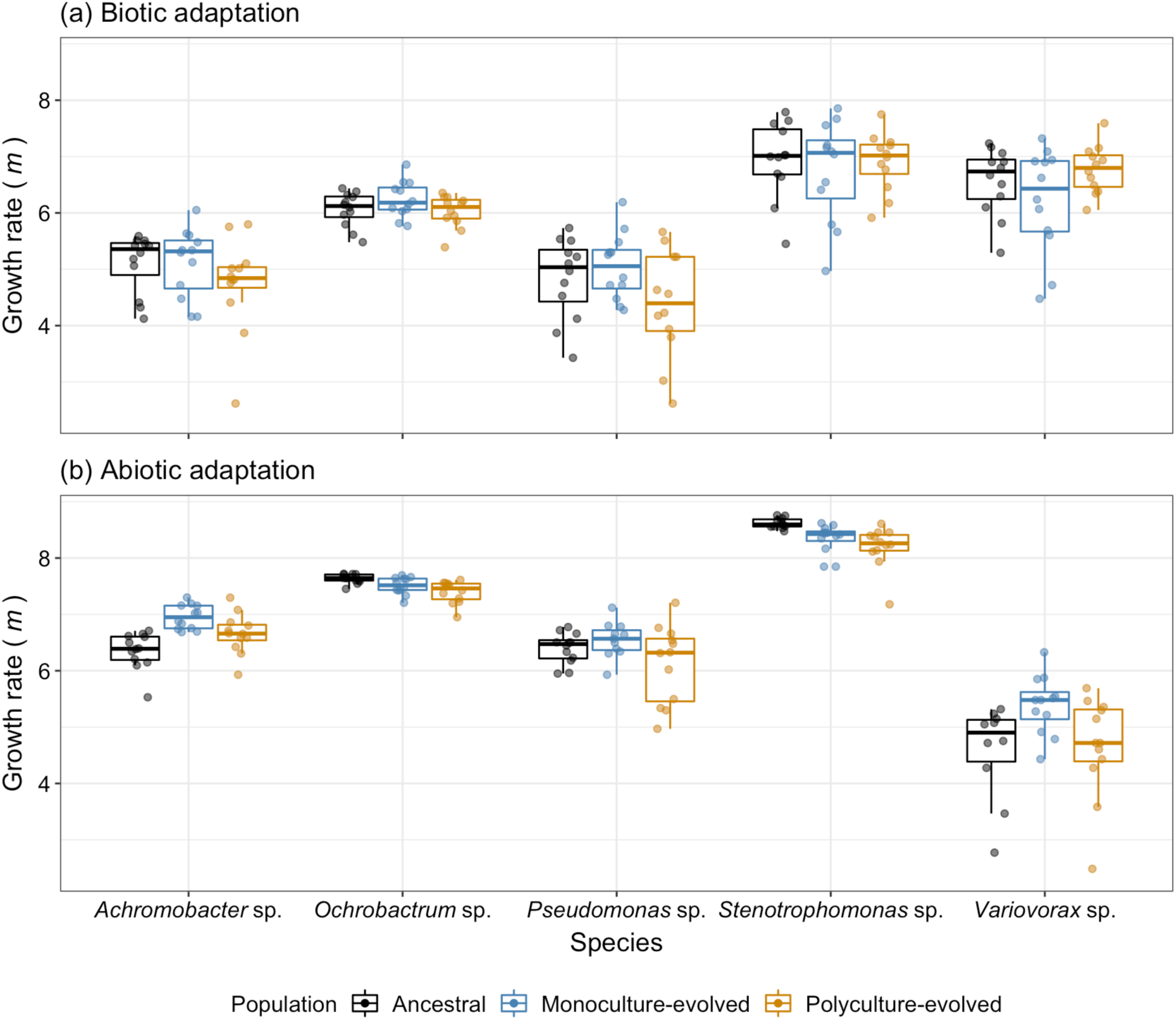
Comparison between growth rates of ancestral, monoculture-evolved and polyculture-evolved populations. (a) Growth rates of focal species were measured within a coevolved community to assess general biotic adaptation. (b) Growth rates were measured in isolation to assess abiotic adaptation. Tops and bottoms of the bars represent the 75th and 25th percentiles of the data, the middle lines are the medians, and the whiskers extend from their respective hinge to the smallest or largest value no further than 1.5* interquartile range.

To estimate abiotic adaptation, we measured growth of the same clones as above in monoculture. The growth rate of polyculture-evolved clones did not differ from their respective ancestors, and only two monoculture-evolved populations showed evidence of abiotic adaptation. While there was a significant interaction between species identity and evolutionary history (F_8, 31.5_ = 3.64, p < 0.001), comparisons between ancestral, monoculture- and polyculture-evolve populations showed that this was driven only by populations evolved in monoculture (Figure 2b). Within species, *Achromobacter* sp. monoculture-evolved clones had the highest growth rate (*m* = 6.95, 95%CI = 6.7 - 7.2) and this was significantly higher than the ancestral growth rate (*m* = 6.35, 95%CI = 6.1 - 6.6; p = 0.003). Growth rates of polyculture-evolved *Achromobacter* sp. clones were intermediate (*m* = 6.66, 95%CI = 6.4 - 6.9), and non-significantly different, to the ancestral (p = 0.204) or monoculture-evolved clones (p = 0.231). Monoculture-evolved *Variovorax* sp. clones had a significantly higher growth rate (*m* = 5.39, 95%CI = 5.1 - 5.6) than the ancestral (*m* = 4.58, 95%CI = 4.3 - 4.9; p < 0.001) and polyculture-evolved clones (*m* = 4.65, 95%CI = 4.4 - 4.9; p < 0.001); and no significant difference in growth rate was evidenced between ancestral and polyculture-evolved *Variovorax* sp. clones (p = 0.934). There were no other significant comparisons between ancestral, monoculture- and polyculture-evolved clones of the other four species (Tukey HSD pairwise comparisons: p > 0.05; Table S2; Figure 2b). Overall, we did not find evidence of significant adaptation to the communities species’ evolved in and abiotic adaptation was only shown by two species evolved in monoculture (Figure 2).

### Local adaptation between (co)evolving communities

To determine the extent of local adaptation to the specific replicate (co)evolving community, we measured the growth rate of each species’ population in the presence of the other four species that had (co)evolved with compared to when growing with four other species from a paired, allopatric community (Figure 1). Growth rates were on average significantly lower in sympatric versus allopatric communities (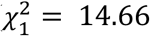, p < 0.001) when measured across all species (interaction between species and sympatric/allopatric community: 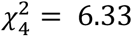, p = 0.177; fixed effect of species 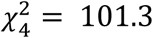, p < 0.001; Figure 3). Despite the lack of significant interaction with species, 11/12 *Pseudomonas* sp. clones had higher growth rates within an allopatric community (mean growth rate difference [m_diff_ = m_allopatric_ - m_sympatric_] = 0.69, ± SE = 0.24); 9/12 *Ochrobactrum* sp. (m_diff_ = 0.34, ± SE = 0.13), 8/12 *Achromobacter* sp. (m_diff_ = 0.74, ± SE = 0.35) and 8/12 *Variovorax* sp. (m_diff_ = 0.30, ± SE = 0.14) clones showed a similar trend (Figure 3). Only 5/12 *Stenotrophomonas* sp. clones had a higher growth rate in allopatric over sympatric communities (m_diff_ = 0.06, ± SE = 0.24). This result strongly suggests that populations were locally maladapted in four out of five species (Table S3; Figure 3).

**Figure 3.**
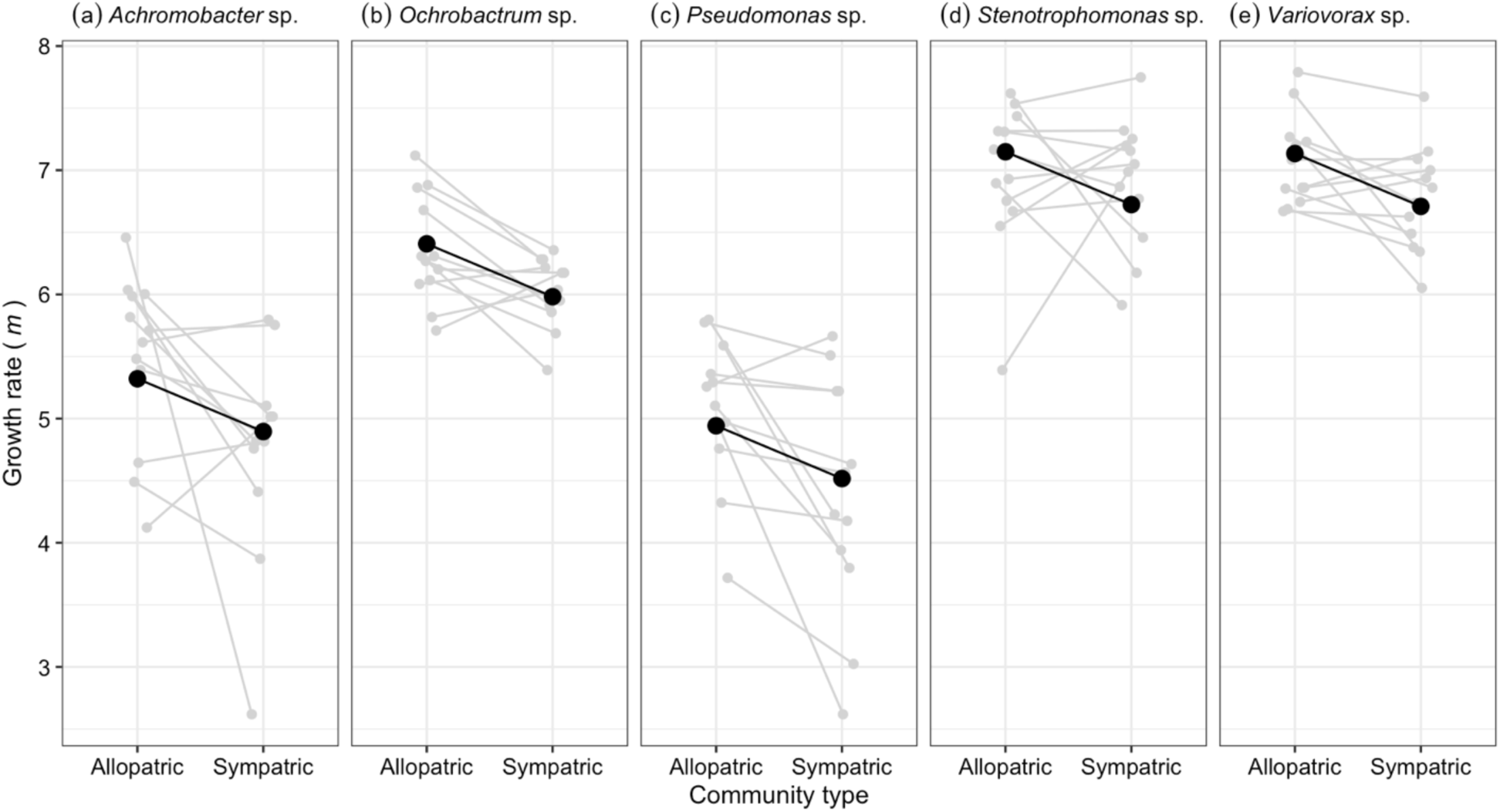
Growth rates (*m*) of species (a - e) in sympatric and allopatric communities. Grey points and lines represent the growth rates of individual clones measured across both community types. Large black points and lines represent predicted growth rates (from mixed effects models) of each species across each community type.

### Community level properties

We have previously shown that all five species can stably coexist (i.e. each can invade from rare) under these culture conditions (Padfield *et al*. bioRxiv). (Co)evolution did not significantly affect this community stability: fitness when rare was significantly greater than 1 for all species (One sample t-test, adjusted p_adj_ < 0.001; Table S4); and relative invader fitness did not differ in communities assembled from the coevolved sympatric populations and communities assembled from monoculture-evolved populations (fixed effect of treatment: F_1,115_ = 2.27, p = 0.135; treatment by species interaction: F_4,114_ = 2.07, p = 0.089; Figure 4).

**Figure 4.**
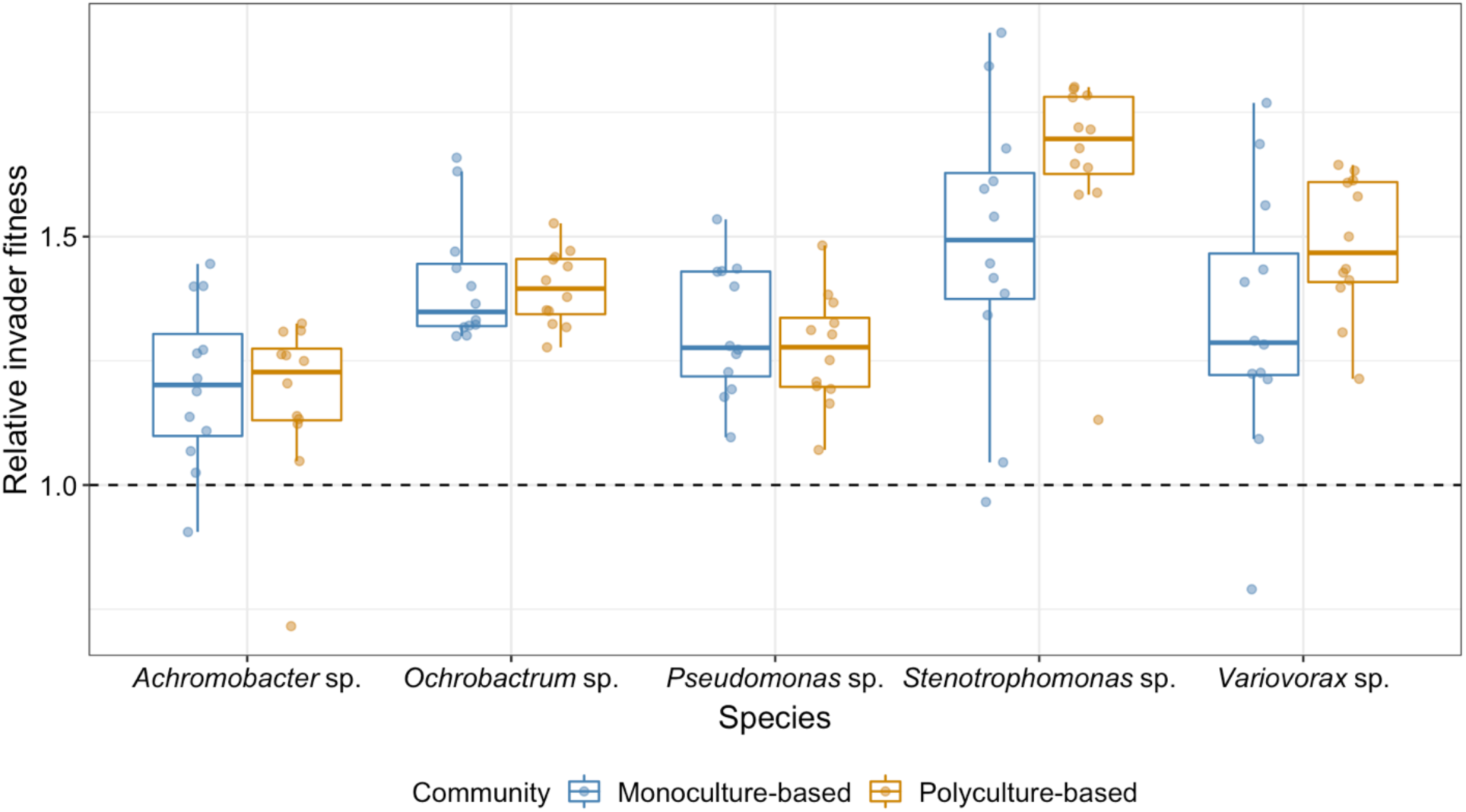
Community stability of monoculture- and polyculture-based communities. An invader fitness greater than 1 indicates successful invasions from rare. Dashed line (y = 1) indicates the relative invader fitness threshold an invader must meet or exceed in order to stably coexist with the other community members. Tops and bottoms of the bars represent the 75th and 25th percentiles of the data, the middle lines are the medians, and the whiskers extend from their respective hinge to the smallest or largest value no further than 1.5* interquartile range.

We also assessed whether evolutionary history impacted community productivity. As local adaptation assays showed species growth rates were greater in allopatric communities relative to sympatric communities, we determined if randomly assembled communities from allopatric polycultures achieved higher total densities than sympatric and monoculture-assembled communities. Overall, there was no significant effect of evolution on productivity between communities composed of monoculture- or polyculture-evolved (sympatric/allopatric) clones (F_2,33_ = 1.9, p = 0.165; Figure 5a). However, the structure (species proportions) and consequently the relative contribution of each species to productivity varied between communities, as apparent from a significant interaction between species identity and community assembly (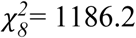, p < 0.001; Figure 5b). Significant pairwise comparisons are summarised in Table S5.

**Figure 5.**
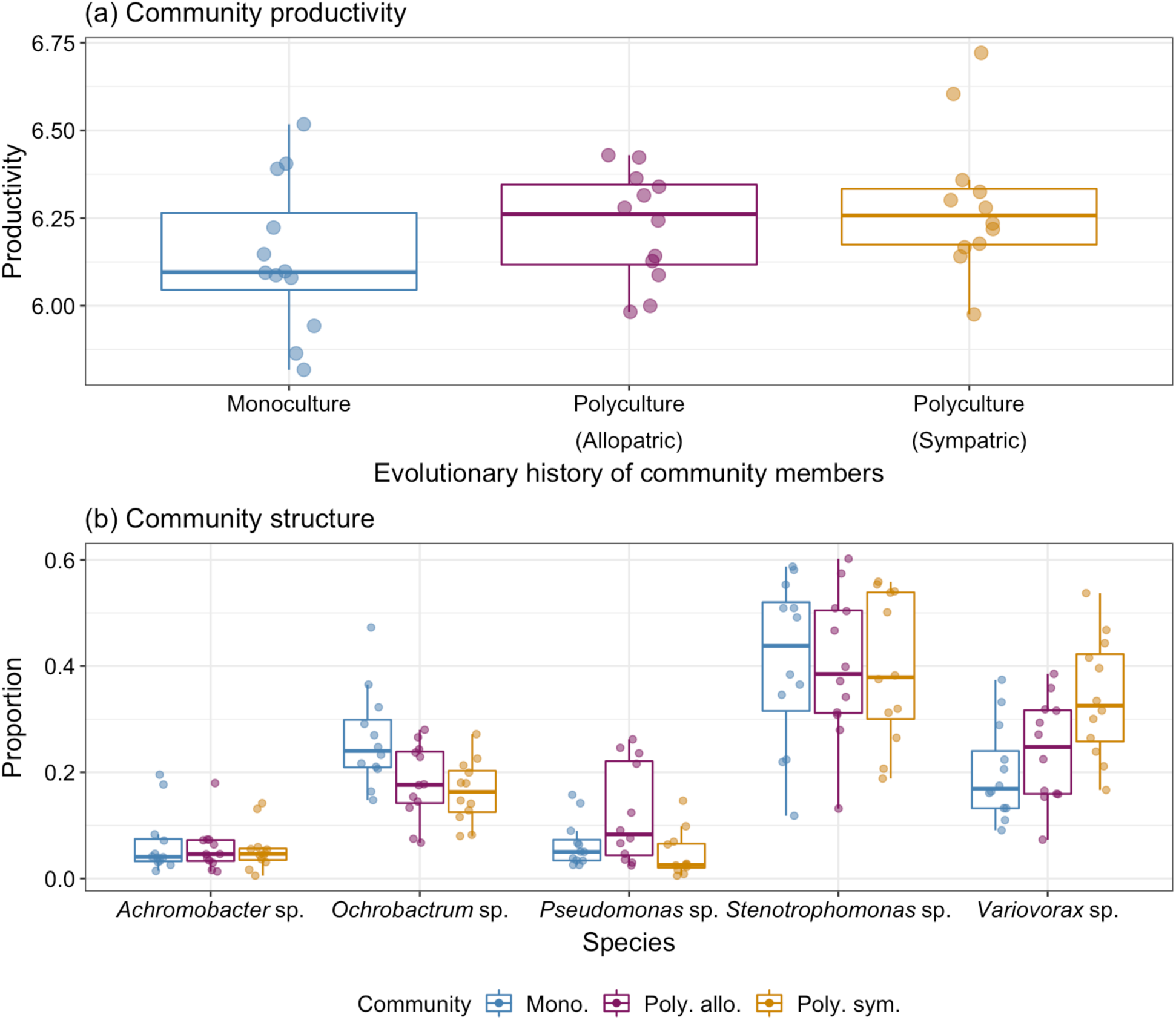
Community productivity and structure of monoculture- and sympatric/allopatric polyculture-evolved communities. (a) Community productivity (change in cellular density (CFUs) over one week) between communities assembled from monoculture- (“Mono.”) and polyculture-evolved (“Poly.”) populations. Polyculture-based communities were assembled from populations evolved in different allopatric (“allo.”) or within the same sympatric (“sym.”) evolution line. (b) The contribution of each species to overall productivity as shown by differing species proportions within the final community. Tops and bottoms of the bars represent the 75^th^ and 25th percentiles of the data, the middle lines are the medians, and the whiskers extend from their respective hinge to the smallest or largest value no further than 1.5* interquartile range.

## Discussion

Here, we investigated patterns of biotic local adaptation of microbial species evolved in multi-species competitive communities for approximately 100 generations. We found that four out of five species typically showed lower growth rates in their own versus foreign communities, suggesting that populations on average were locally maladapted. This local maladaptation did not affect the stability of the community (all species could still invade from rare (Chesson 2018) equally well in sympatric polyculture and monoculture-based communities) or productivity. Moreover, despite this clear evidence of rapid evolutionary change, we found no evidence that populations evolved in polyculture adapted to the biotic environment *per se*, and significant abiotic adaptation was only detected in species evolved in monoculture.

We do not know the mechanism underpinning local (mal)adaptation and attempting to uncover it is beyond the scope of this study. We speculate that rather than species showing character displacement to use different resources to escape competition (Pacala & Roughgarden 1982), they instead show phenotypic convergence which increases competition (Abrams 1987). Evolutionary convergence in life history traits in response to co-culturing has recently been shown in experimentally evolved populations of duckweed (Hart *et al*. 2019). Theoretically, such convergence can arise when species compete for non-substitutional resources, whereby more efficient use in one species drives enhanced use in competitors (Abrams 1987; Fox & Vasseur 2008). If the evolved responses differ between communities, this could result in species on average performing worse in their own community than in other communities owing to greater niche overlap in the former. It is possible that competitors undergo divergent antagonistic coevolution between communities by a mechanism (Zhang *et al*. 2009) such as production of, and defence against, anti-competitor toxins (Inglis *et al*. 2016; Granato *et al*. 2019). However, this hypothesis seems less likely because total densities, which are expected to be reduced by anti-competitor toxins, did not differ between allopatric and sympatric community combinations.

Contrary to some previous research (Lawrence *et al*. 2012; Gómez & Buckling 2013; Briscoe Runquist *et al*. 2020), we did not observe any net growth improvements in the presence of the community for community-evolved populations i.e. monoculture-, polyculture-evolved and ancestral populations all had the same growth rate. Taken together with the broadly consistent pattern of local maladaptation of populations to their sympatric communities, this finding is consistent with antagonistic “red queen” coevolutionary dynamics between competitors, whereby an adaptation of one population is offset by a counter-adaptation in others (Valen 1973; Stenseth & Smith 1984). However, detailed measurement of coevolutionary dynamics between several species (Buckling & Rainey 2002; Hall *et al*. 2020) would be necessary to unequivocally demonstrate this. Consistent with previous work (Hereford 2009; Lawrence *et al*. 2012; Gómez & Buckling 2013; Hall *et al*. 2018; Briscoe Runquist *et al*. 2020), adaptation of at least one species to the abiotic environment was constrained. Specifically, *Variovorax* sp. monoculture-evolved clones had a higher growth rate than both the ancestral and polyculture-evolved clones when growing in isolation. It is unclear why the effect was strongest for *Variovorax* sp. but it may be because this is the only species that appears to benefit from the presence of other species (Padfield *et al*., bioRxiv), hence there may have been strong selection to use alternative resources when evolved in monoculture.

There has been rapidly growing interest in experimental evolution of multispecies communities (Lawrence *et al*. 2012; Celiker & Gore 2014; Pantel *et al*. 2015; Ghoul & Mitri 2016; Hall *et al*. 2018; terHorst *et al*. 2018) because such studies may have greater relevance to nature than do monoculture experiments. However, these laboratory-based model systems are still far removed from natural systems and, in combination with limited theoretical predictions, it is unclear how far results from these studies can be generalised. Nevertheless, one advantage of our system is that it captures a key feature of natural communities: the species can stably coexist (Chesson 2018; Padfield *et al*. bioRxiv). This is very rarely demonstrated in other model communities and may often not be the case.

The rapid evolution of local (mal)adaptation has potentially significant implications for microbial community structure and function in environmental, medical and biotechnology contexts; over and above an important role for rapid evolution in community ecology in general (Johnson & Stinchcombe 2007; Widder *et al*. 2016). Local maladaptation will increase the invasion success of both single species (Richardson & Pyšek 2007) and entire communities (Castledine *et al*. 2020), with the latter in particular potentially causing rapid changes in community composition and function (Sierocinski *et al*. 2017; Wang *et al*. 2019). Moreover, while we not observe detectable (negative) changes in measures of community function, this is likely to be an outcome if the extent of local maladaptation increases through time.

## Data accessibility statement

All data and R code used in the analysis will be made available on GitHub.

## Acknowledgements

We thank Matthew Silk for helpful advice on the statistical analysis. This work was funded by NERC.

