## Supplementary Information for "Experimental multi-species microbial (co)evolution results in local maladaptation"

Table S1: Replicates removed due to contamination

Table S2: Multiple pairwise comparisons of growth rates in isolation (abiotic adaptation)

Table S3: Parameter estimates and 95%CI of growth rates in sympatry and allopatry

Table S4. Model parameters for one-sample t-tests of each species' relative invader fitness

Table S5: Multiple pairwise comparisons of species proportions in communities

Table S1. Replicates removed due to contamination. Unless otherwise stated, no replicates were removed from unmentioned datasets, thus retaining a total of 12 replicates per species and treatment.

| Dataset | Species | Evolutionary history | N° replicates removed |
|---|---|---|---|
| Abiotic adaptation | S | Ancestor | 1 |
| | P | Monoculture | 1 |
| | V | Ancestor | 2 |

Table S2. Multiple pairwise comparisons comparing growth rates (*m*) between ancestral ("Anc", monoculture- ("Mono.") and polyculture-adapted ("Poly.") populations when in isolation. Degrees of freedom ("df") calculated using the Kenward-Roger method and p-values adjusted using the Tukey method of comparing a family of four estimates. Significant contrasts in bold.

| Contrast | Sp. | Estimate | SE | df | t-ratio | p-value |
|---|---|---|---|---|---|---|
| **Anc. - Mono.** | **A** | **-0.605** | **0.181** | **161** | **-3.353** | **0.003** |
| Anc. - Poly. | A | -0.309 | 0.181 | 161 | -1.711 | 0.204 |
| Mono. - Poly. | A | 0.297 | 0.181 | 161 | 1.642 | 0.231 |
| Anc. - Mono. | O | 0.122 | 0.181 | 161 | 0.677 | 0.777 |
| Anc. - Poly. | O | 0.238 | 0.181 | 161 | 1.319 | 0.386 |
| Mono. - Poly. | O | 0.116 | 0.181 | 161 | 0.643 | 0.797 |
| Anc. - Mono. | P | -0.151 | 0.185 | 161 | -0.817 | 0.693 |
| Anc. - Poly. | P | 0.28 | 0.181 | 161 | 1.549 | 0.271 |
| Mono. - Poly. | P | 0.43 | 0.185 | 161 | 2.332 | 0.054 |
| Anc. - Mono. | S | 0.28 | 0.185 | 161 | 1.514 | 0.287 |
| Anc. - Poly. | S | 0.419 | 0.185 | 161 | 2.272 | 0.063 |
| Mono. - Poly. | S | 0.14 | 0.181 | 161 | 0.775 | 0.719 |
| **Anc. - Mono.** | **V** | **-0.81** | **0.189** | **161** | **-4.279** | **<0.001** |
| Anc. - Poly. | V | -0.066 | 0.189 | 161 | -0.351 | 0.934 |
| **Mono. - Poly.** | **V** | **0.744** | **0.181** | **161** | **4.119** | **<0.001** |

Table S3. Model parameters values of species' ("Sp.") growth rates in allopatric and sympatric communities with standard error ("SE") and 95% confidence intervals ("CI"). Degrees of freedom ("df") calculated using the Kenward-Roger method. Effect of community type significant at p = 0.0002 across all species.

| Community type | Sp. | Mean | SE | df | Lower CI | Upper CI |
|---|---|---|---|---|---|---|
| Allopatric | A | 5.321 | 0.141 | 72.637 | 5.039 | 5.603 |
| Sympatric | A | 4.895 | 0.141 | 72.637 | 4.614 | 5.177 |
| Allopatric | O | 6.407 | 0.141 | 72.637 | 6.125 | 6.689 |
| Sympatric | O | 5.982 | 0.141 | 72.637 | 5.7 | 6.264 |
| Allopatric | P | 4.942 | 0.141 | 72.637 | 4.66 | 5.224 |
| Sympatric | P | 4.517 | 0.141 | 72.637 | 4.235 | 4.799 |
| Allopatric | S | 7.149 | 0.141 | 72.637 | 6.867 | 7.431 |
| Sympatric | S | 6.723 | 0.141 | 72.637 | 6.441 | 7.005 |
| Allopatric | V | 7.134 | 0.141 | 72.637 | 6.853 | 7.416 |
| Sympatric | V | 6.709 | 0.141 | 72.637 | 6.427 | 6.991 |

Table S4. Model parameters for multiple one-sample t-tests testing the relative invader fitness of each species against 1 within each community type. P-values adjusted ("Adj.") using the false discovery rate method.

| Sp. | Community type | Estimate | t-value | p-value | df | Lower CI | Upper CI | Adj. p-value |
|---|---|---|---|---|---|---|---|---|
| A | Polyculture-based | 1.173 | 3.562 | 0.004 | 11.000 | 1.066 | 1.281 | 0.004 |
| A | Monoculture-based | 1.202 | 4.27 | 0.001 | 11.000 | 1.098 | 1.307 | 0.001 |
| O | Polyculture-based | 1.397 | 18.31 | <0.001 | 11.000 | 1.349 | 1.444 | <0.001 |
| O | Monoculture-based | 1.405 | 11.237 | <0.001 | 11.000 | 1.325 | 1.484 | <0.001 |
| P | Polyculture-based | 1.272 | 8.363 | <0.001 | 11.000 | 1.2 | 1.343 | <0.001 |
| P | Monoculture-based | 1.312 | 8.183 | <0.001 | 11.000 | 1.228 | 1.395 | <0.001 |
| S | Polyculture-based | 1.655 | 12.438 | <0.001 | 11.000 | 1.539 | 1.771 | <0.001 |
| S | Monoculture-based | 1.482 | 5.911 | <0.001 | 11.000 | 1.302 | 1.661 | <0.001 |
| V | Polyculture-based | 1.481 | 12.002 | <0.001 | 11.000 | 1.393 | 1.569 | <0.001 |
| V | Monoculture-based | 1.332 | 4.319 | 0.001 | 11.000 | 1.163 | 1.501 | 0.001 |

Table S5. Multiple pairwise comparisons comparing community structure of polyculture-based ("Poly.") communities, in which populations had evolved in sympatry ("sym.") or allopatry ("allo."), and monoculture- based ("Mono.") communities. In a general linear model, species' proportions were tested against fixed effects of community type and species ("Sp.") with a quasibinomial error structure. Tests performed on the log odds ratio scale. Significant contrasts in bold.

| Contrast | Sp. | Odds-ratio | SE | z-ratio | p-value |
|---|---|---|---|---|---|
| Mono. / Poly. allo. | A | 1.131 | 0.454 | 0.307 | 0.949 |
| Mono. / Poly. sym. | A | 1.216 | 0.487 | 0.489 | 0.876 |
| Poly. allo. / Poly. sym. | A | 1.076 | 0.417 | 0.188 | 0.981 |
| Mono. / Poly. allo. | O | 1.518 | 0.349 | 1.816 | 0.164 |
| **Mono. / Poly. sym.** | **O** | **1.917** | **0.449** | **2.774** | **0.015** |
| Poly. allo. / Poly. sym. | O | 1.263 | 0.296 | 0.995 | 0.58 |
| Mono. / Poly. allo. | P | 0.526 | 0.186 | -1.818 | 0.164 |
| Mono. / Poly. sym. | P | 1.578 | 0.669 | 1.074 | 0.53 |
| **Poly. allo. / Poly. sym.** | **P** | **3.001** | **1.1** | **2.997** | **0.008** |
| Mono. / Poly. allo. | S | 1.083 | 0.211 | 0.409 | 0.912 |
| Mono. / Poly. sym. | S | 1.047 | 0.2 | 0.241 | 0.968 |
| Poly. allo. / Poly. sym. | S | 0.967 | 0.174 | -0.186 | 0.981 |
| Mono. / Poly. allo. | V | 0.757 | 0.177 | -1.195 | 0.456 |
| **Mono. / Poly. sym.** | **V** | **0.46** | **0.102** | **-3.5** | **0.001** |
| **Poly. allo. / Poly. sym.** | **V** | **0.608** | **0.12** | **-2.528** | **0.031** |
